# Estimating the current burden of Chagas disease in Mexico: a systematic review of epidemiological surveys from 2006 to 2017

**DOI:** 10.1101/423145

**Authors:** Audrey Arnal, Etienne Waleckx, Claudia Herrera, Eric Dumonteil

**Affiliations:** Centro de Investigaciones Regionales Dr Hideyo Noguchi, Universidad Autónoma de Yucatán, calle 96 s/n x av. Jacinto Canek y calle 47, Col. Paseo de las Fuentes, CP 97225, Mérida, Yucatán, México; Department of Tropical Medicine, School of Public Health and Tropical Medicine, and Vector-Borne and Infectious Disease Research Center, Tulane University, 1440 Canal St., New Orleans, La 70112, USA

## Abstract

**Background:** In Mexico, estimates of Chagas disease prevalence and burden vary widely. Updating surveillance data is therefore an important priority to ensure that Chagas disease does not remain a barrier to the development of Mexico’s most vulnerable populations.

**Methodology/Principal Findings:** The aim of this systematic review was to analyze the literature on epidemiological surveys to estimate Chagas disease prevalence and burden in Mexico, during the period 2006 to 2017. A total of 2,764 articles were screened and 38 were retained for the final analysis. Epidemiological surveys have been performed in most of Mexico, but with variable geographic coverage. Based on studies reporting confirmed cases, i.e. using at least 2 serological tests, the national seroprevalence of *Trypanosoma cruzi* infection was 2.26% [95% Confidence Interval (CI) 2.12-2.41], suggesting that there are 2.71 million cases in Mexico. Studies focused on pregnant women, which may transmit the parasite to their newborn during pregnancy, reported a seroprevalence of 1.00% [95% CI 0.87-1.14], suggesting that there are 22,930 births from *T. cruzi* infected pregnant women per year, and 1,445 cases of congenitally infected newborns per year. Children under 18 years had a seropositivity rate of 1.49% [95% CI 1.20-1.85], which indicate ongoing transmission. Finally, cases of *T. cruzi* infection in blood donors have also been reported in most states, with a national seroprevalence of 0.51% [95% CI 0.49-0.53].

**Conclusions/Significance:** Our analysis suggests a disease burden for *T. cruzi* infection higher than previously recognized, highlighting the urgency of establishing Chagas disease surveillance and control as a key national public health priority in Mexico, to ensure that it does not remain a major barrier to the economic and social development of the country’s most vulnerable populations.

****Author summary**:** In Mexico, estimates of Chagas disease prevalence and burden vary widely due to the ecology and epidemiology of this disease resulting of many geographical, ecological, biological, and social interactions. Better data are thus urgently needed to help develop appropriate public health programs for disease control and patient care. In this study we analyzed published data on *T. cruzi* seroprevalence infection in Mexico between 2006 and 2017. This systematic review shows a national seroprevalence of *T. cruzi* infection of 2.26% [95%CI 2.12-2.41], with over 2.71 million cases in Mexico, which is higher than previously recognized. The presence of *T. cruzi* infection in specific subpopulations such as pregnant women, children and blood donors also informs on specific risks of infection and call for the implementation of well-established control interventions. This work confirms the place of Mexico as the country with the largest number of cases, highlighting the urgency of establishing Chagas disease control as a key national public health priority.

## Introduction

Chagas disease or American trypanosomiasis is an infection caused by the protozoan parasite *Trypanosoma cruzi*, which is mainly transmitted to humans and other mammals through the contaminated feces of hematophagous bugs called triatomines (family Reduviidae). However, it can also be spread via non-vectorial routes, such as blood transfusion, congenital transmission, organ transplantation, ingestion of food and beverages contaminated with *T. cruzi* or laboratory accidents [1]. Over the years, infection with *T. cruzi* can cause heart failure or sudden death associated with progressive heart damage [2]. Some patients may also suffer from digestive, neurological or multiple alterations. This disease, classified by the World Health Organization (WHO) within the group of neglected tropical diseases, is a major public health problem in Latin America where it is estimated that 6 to 7 million people are currently infected [1]. Due to human migrations, Chagas disease is emerging in other regions (Europe and United States principally) [3]. Estimates suggest that 80,000 to 120,000 *T. cruzi*-infected immigrants live in Europe, and 300,000 live in the United States [4], and the disease is a growing concern in these regions [5]. The global economic burden of Chagas disease is more than US$ 7.2 billion per year, exceeding the costs of other diseases of health impact such as certain cancer (US$6.7 billion for uterine cancer, US$4.7 billion for cervical cancer, and US$5.3 billion for oral cancer) or rotavirus infections (US$ 2 billion) [6,7].

In Mexico, estimates of Chagas disease prevalence and burden vary widely, which has complicated the establishment of a strong National Chagas Disease Program for vector control as well as for patient detection and care in the country. For the past several years, the Ministry of Health only reports a few hundred cases per year [8], suggesting that the disease has an anecdotal burden in terms of public health. On the other hand, other estimates suggest that there are about 1.1 million individuals infected with *T. cruzi* in Mexico, and 29.5 million at risk of infection [9,10]. Higher estimates of up to 6 million cases have also been proposed [11]. The annual cost for medical care for patients in the outpatient setting in this country is estimated between US$4,463 and US$9,601, and annual costs for patients admitted via an emergency care unit is between US$6,700 and US$11,838 [12].

There are also important regional differences in prevalence levels or number of cases reported in Mexico. For example, between 1928 and 2004 the states with the highest number of human cases reported were Chiapas, Guerrero, Jalisco, Morelos, Nayarit, Oaxaca and Queretaro. Conversely, few cases were reported in the states of Chihuahua, Coahuila, Guanajuato and Estado de Mexico [11]. It is not clear if such differences in prevalence are reflecting true differences in eco-epidemiological conditions, as Mexico is home to an extensive diversity of triatomine species, habitats, and socioeconomic conditions, or if there are bias in disease surveillance among regions [11].

Such wide discrepancies are important to reconcile to ensure that Chagas disease does not remain a major barrier to the development of Mexico’s most vulnerable populations. Updating and improving surveillance data for Chagas disease in Mexico is therefore an important public health priority. In this context, the aim of this systematic review was to analyze available literature on epidemiological surveys to estimate Chagas disease prevalence and burden in Mexico. We focused our study on the period from 2006 to 2017, to define current disease status rather than historical/cumulative burden, but our results are nonetheless compared with past reviews [11,13,14] to shed light on possible temporal trends on the status of Chagas disease in the country.

## Methods

This systematic review was conducted in accordance with the PRISMA statement [15] (Supporting information). Potential data sources were identified and selected in different bibliographic databases. The ISI Web of Science (v5.13.1) was chosen because it incorporates many relevant databases including the SciELO Citation Index from 1997 onwards (provides access to leading journals from Latin America, Portugal, Spain and South Africa) and the Web of Science’s Core Collection from 1980 onwards (https://webofknowledge.com/). A part of the literature was selected from the LILACS database (lilacs.bvsalud.org/en/), which is the most important index of scientific and technical literature of Latin America and the Caribbean. Finally, the BibTri database (https://bibtri.cepave.edu.ar/) was also used because it integrates scientific literature specifically related to Chagas disease.

We restricted our search to the period from January 2006 to December 2017, to obtain information on the current status of Chagas disease in Mexico rather than on its historical/cumulative status, which has been summarized in previous reviews [11,13,14]. Selection was made using the search terms ‘Chagas disease in Mexico/Enfermedad de Chagas en México’.

For all these articles, titles and abstracts were screened for any indication that the study contained data related to the seroprevalence of *T. cruzi* infection in human populations from Mexico. Typically, this excluded studies of, for example, therapeutic options for patients with chronic Chagas disease, molecular studies of lab strains of the parasite, or experimental model developments (Figure 1). In the second step of the process, full text copies were obtained and articles containing quantitative data on *T. cruzi* infection seroprevalence were retained. Extreme care was taken in cross-validating whether the information contained in each study was unique and not duplicated elsewhere.

**Figure 1:**
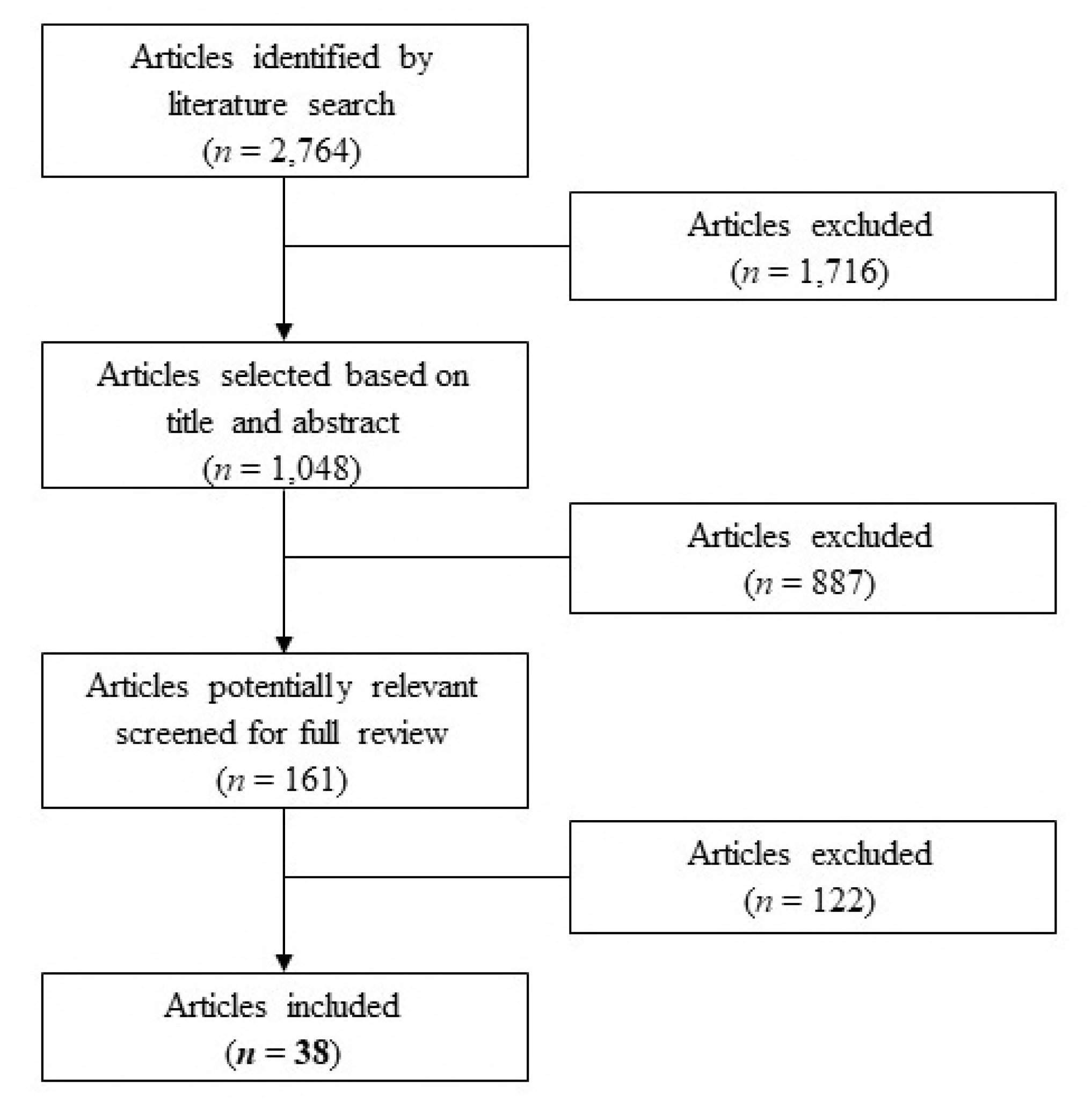
Process flow chart for the identification, screening, eligibility, and inclusion of studies.

The ultimate step was to extract the relevant information contained in the selected articles which included 1) publication data (bibliographic information), 2) sampling dates, 3) sampling strategy (archive, random, volunteers, etc…), 4) geographic area covered by the study, 5) studied population (blood donors, patients, pregnant women, newborns, children, random populations), 6) laboratory techniques used (ELISA tests; IHA, PCR…) and their validation, 7) total sample size, number of human cases, and reported prevalence. If no prevalence was reported, it was calculated from the total of sample size and the number of human cases of *T. cruzi* seropositivity. We further calculated 95% confidence intervals (95%CI) based on the reported data and sample sizes [16,17]. The prevalences obtained were compared to the data reported by the Ministry of Health and with other reviews [8]. Studied populations were divided into subgroups to allow for analysis of the seroprevalence of *T. cruzi* infection at different levels, including the general population, pregnant women, children, and blood donors.

## Results

A total of 2,764 articles were screened and 38 were retained for the final analysis (see Figure 1). 1) All the articles included in this study corresponded to serological surveys in different populations and settings, including general or specific populations such as pregnant women or blood donors, published between 2006 and 2017 (Table 1). Research on Chagas disease seroprevalence has been performed in most of the Mexican Republic (Figure 2), although the number and scale of the studies varied greatly. The states with more studies were Veracruz, Yucatan, and Queretaro.

**Table 1:**
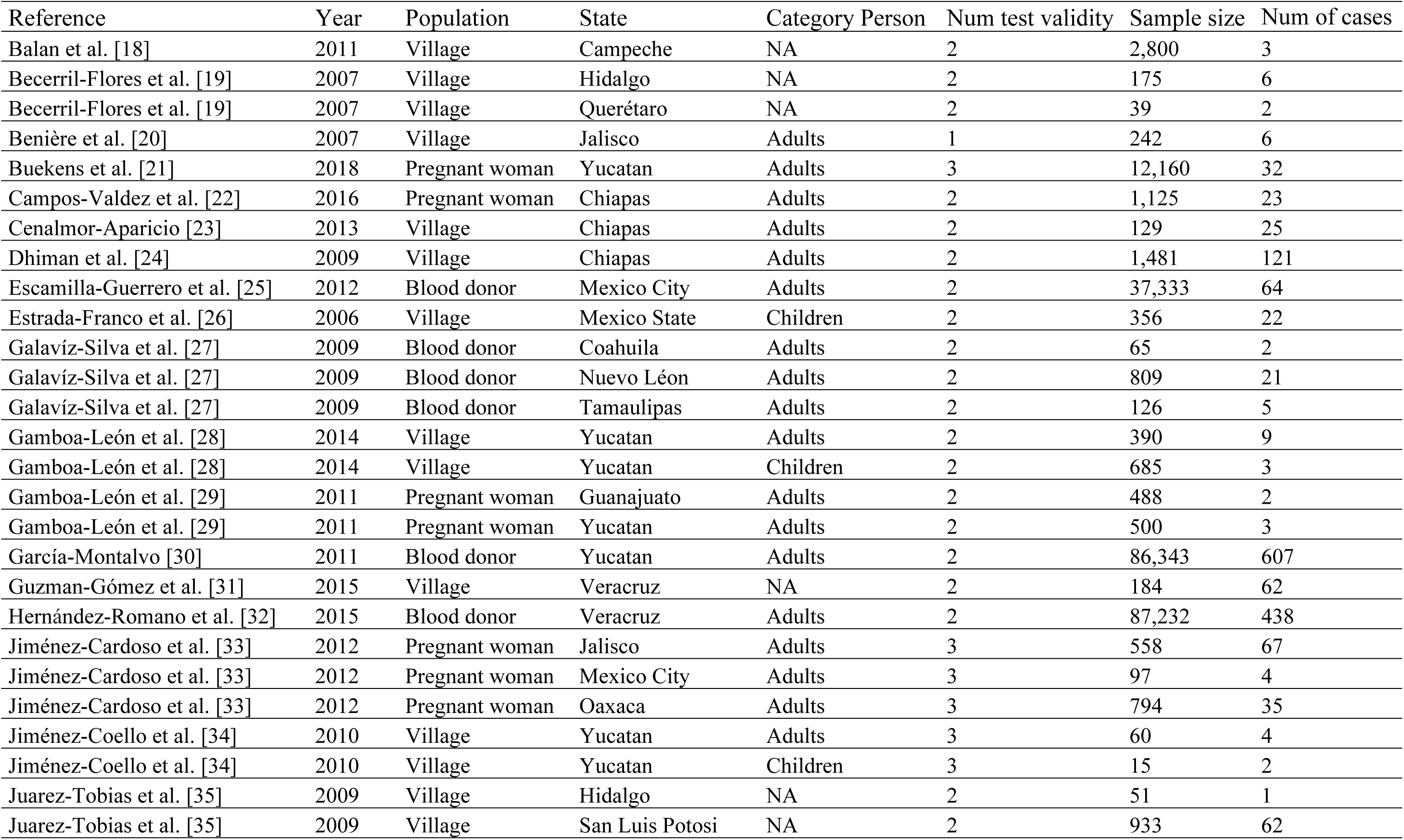
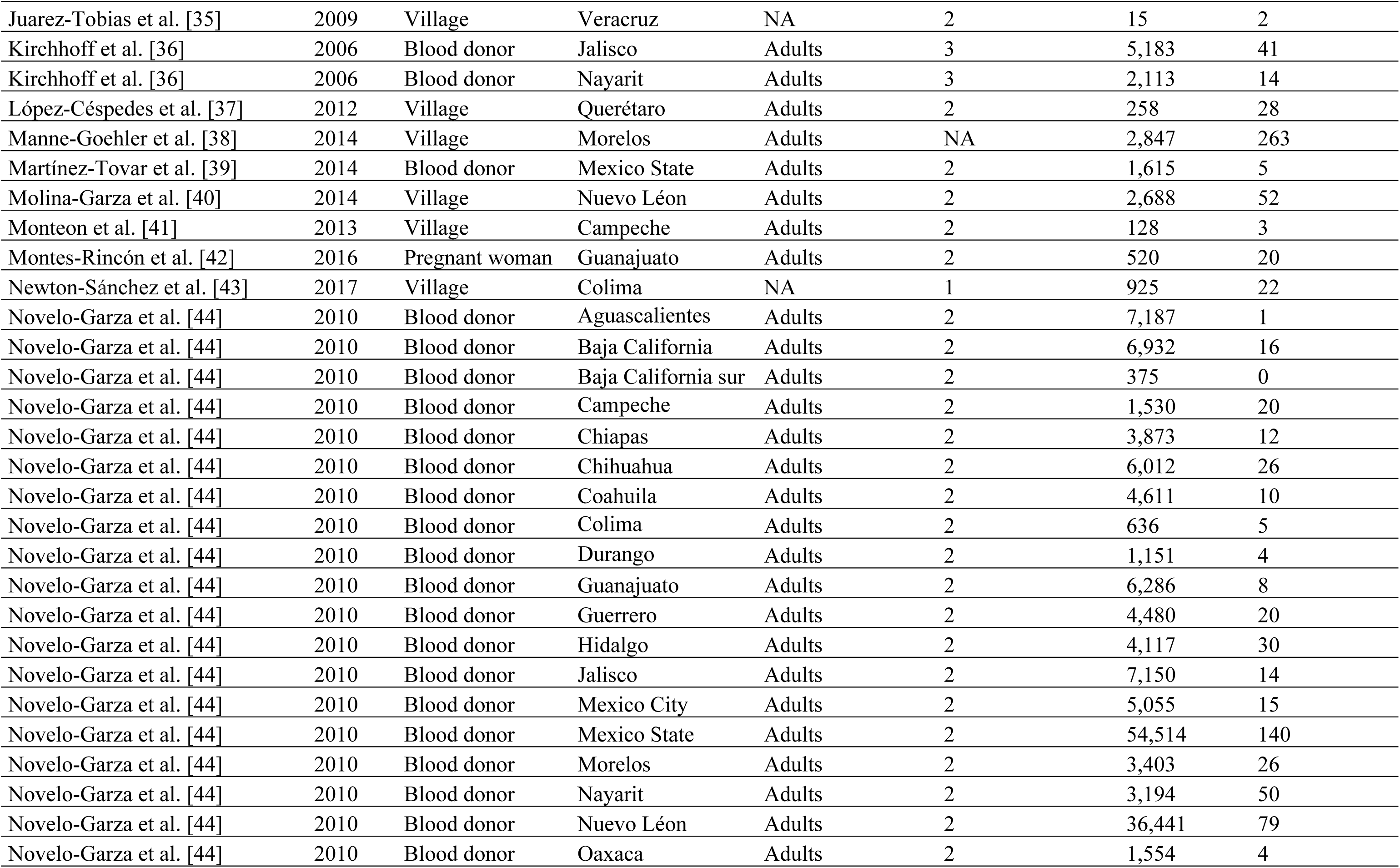
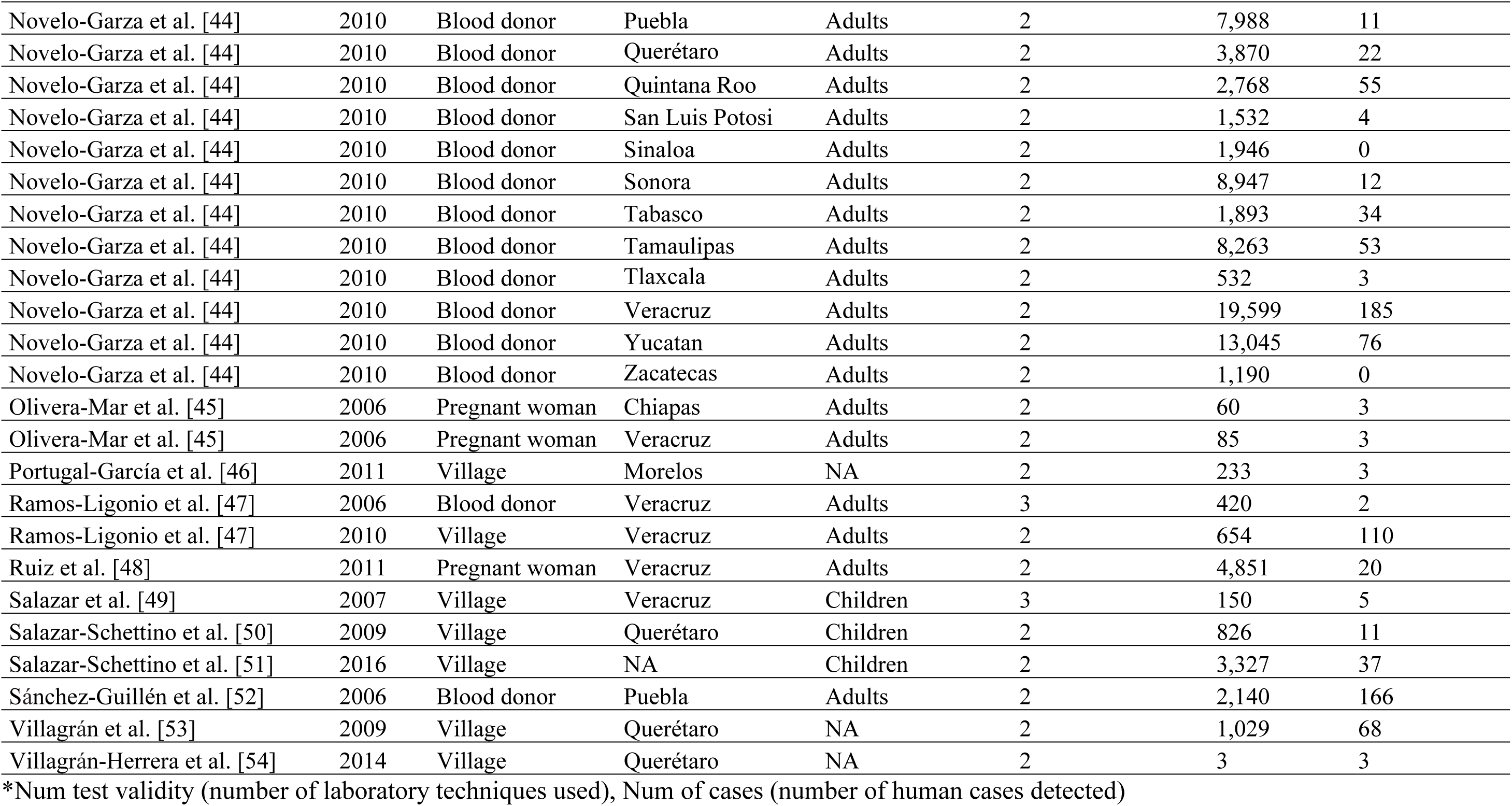
Characteristics of included studies.

**Figure 2:**
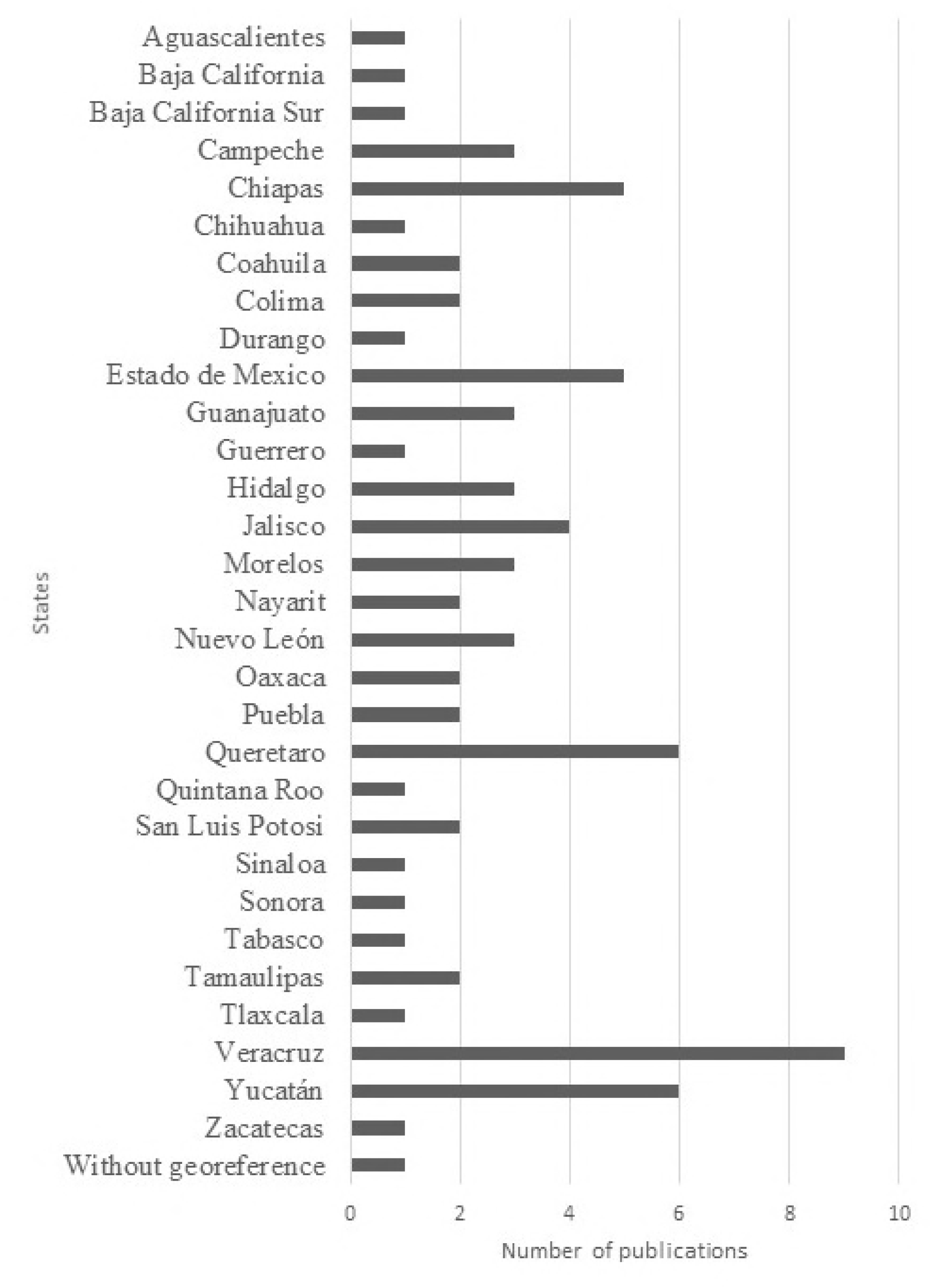
Number of publications which report human cases of *T. cruzi* seropositivity from states of Mexico, 2006-2017. Each publication can cover several states.

We first considered the studies in random populations and in pregnant women (which can be considered as highly representative of the general population as well [55]). When considering all studies, irrespective of the inclusion of confirmatory diagnostic (i.e. based on a single serological test, 29 studies), the total number of human cases reported in the literature during the period 2006-2017 was 1,147 with a national prevalence of 2.74% [95%CI 2.59-2.90] in the general population (Table 2). Seroprevalence of infection varied between 0.21% and 9.13% depending on the state. Only 3 studies were based on a single test and when considering only the studies in which at least 2 serological tests had been performed (26 studies), hence cases had been confirmed as currently recommended by the WHO for an accurate identification of cases, the national seroprevalence was 2.26% [95%CI 2.12-2.41], with seroprevalences between 0.21% and 12.01% depending on the state (Table 3). The highest seroprevalence levels were reported in the states of Jalisco, San Luis Potosi, Chiapas, Estado de Mexico, Queretaro, and Oaxaca. Based on a national population of nearly 120 million (National census of 2015), this seroprevalence level would correspond to 2.71 million cases in the country [95%CI 1.55-3.20 million]. On the other hand, the number of cases of *T. cruzi* infection reported by the national program of epidemiologic surveillance of the Ministry of Health during 2006-2017 period reached 8,687 ([8] and Table 4), with a regular increase in the number of cases detected with time.

**Table 2:** Cases of *Trypanososma cruzi* infection detected in serological surveys of general populations and pregnant women during 2006-2017 (29 studies).

| State | Sample | Cases* | Prevalence (%) | Confidence Interval (95%) |
| --- | --- | --- | --- | --- |
| Campeche | 2,928 | 6 | 0.21 | 0.09-0.44 |
| Chiapas | 2,795 | 172 | 6.15 | 5.32-7.10 |
| Colima | 925 | 22 | 2.37 | 1.58-3.58 |
| Estado de México | 453 | 26 | 5.74 | 3.95-8.28 |
| Guanajuato | 1008 | 22 | 2.18 | 1.44-3.28 |
| Hidalgo | 226 | 7 | 3.10 | 1.51-6.26 |
| Jalisco | 800 | 73 | 9.13 | 7.32-11.33 |
| Morelos | 3,080 | 266 | 8.64 | 7.70-9.68 |
| Nuevo León | 2,688 | 52 | 1.93 | 1.47-2.52 |
| Oaxaca | 794 | 35 | 4.41 | 3.19-6.07 |
| Querétaro | 2,155 | 112 | 5.20 | 4.34-6.22 |
| San Luis Potosí | 933 | 62 | 6.65 | 5.22-8.43 |
| Veracruz | 5,939 | 202 | 3.40 | 2.97-3.89 |
| Yucatán | 13,810 | 53 | 0.38 | 0.29-0.50 |
| Sub Total | 38,534 | 1,110 | 2.88 | 2.72-3.05 |
| Without georef. | 3,327 | 37 | 1.11 | 0.81-1.53 |
| Total | 41,861 | 1,147 | 2.74 | 2.59-2.90 |

\*Cases are reported using 1 or more serological tests to determine the infection.

**Table 3:**
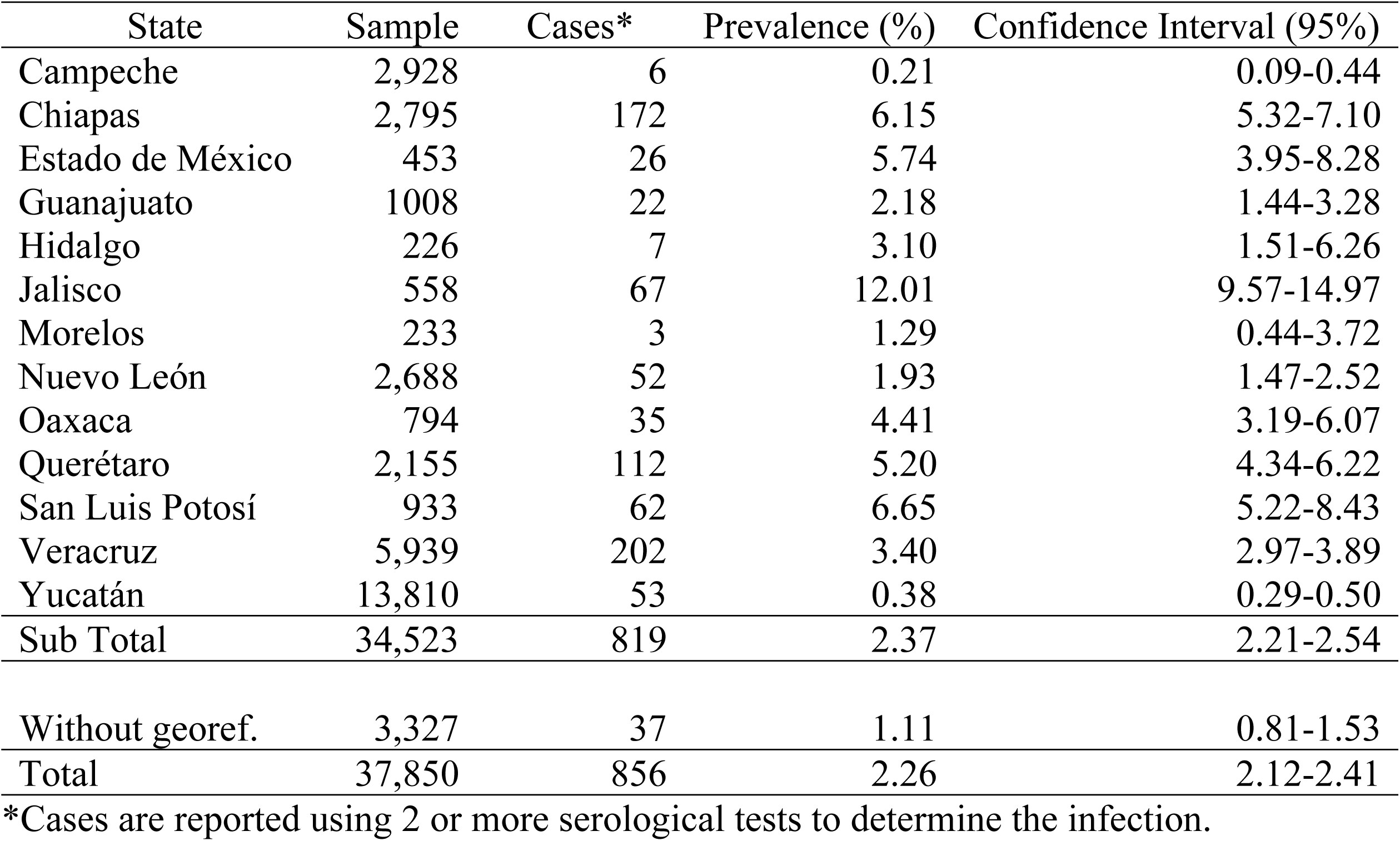
Confirmed cases of *Trypanososma cruzi* infection detected in serological surveys of general populations and pregnant women during 2006-2017 (26 studies).

**Table 4:** Number of new cases of Chagas disease per years, reported by the Ministry of Health, in all the states of Mexico [8].

| Years | Sample |
| --- | --- |
| 2006 | 400 |
| 2007 | 392 |
| 2008 | 679 |
| 2009 | 613 |
| 2010 | 528 |
| 2011 | 801 |
| 2012 | 830 |
| 2013 | 762 |
| 2014 | 735 |
| 2015 | 1,095 |
| 2016 | 994 |
| 2017 | 856 |
| Total | 8,687 |

A few of these studies (7 studies) focused on pregnant women, which may transmit the parasite to their newborn during pregnancy [56]. While these studies only covered 7 states (Table 5), a total of 212 *T. cruzi*-infected pregnant women were detected, for a global seroprevalence of *T. cruzi* infection of 1.00% [95%CI 0.87-1.14] in this specific population. The highest seroprevalence levels in pregnant women were reported in the states of Jalisco, Oaxaca, and Estado de Mexico. Based on current birth rate in Mexico (2,293,000 births in 2016), this would correspond to 22,930 births from *T. cruzi* infected pregnant women per year. With a congenital transmission rate of 6.3% [21], there may be 1,445 cases of congenitally infected newborns per year in the country.

**Table 5:**
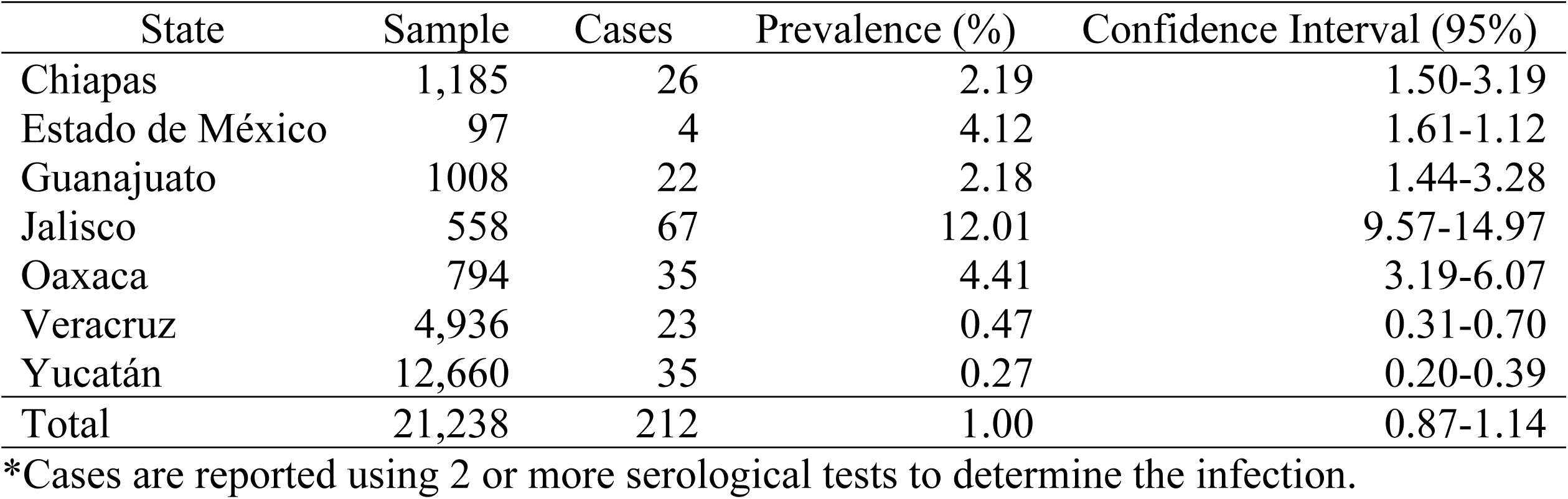
Prevalence of positive pregnant women by state, 2006-2017 (7 studies).

Some studies also focused on or included data on children under 18 years (6 studies), which may indicate more recent transmission. These covered only 4 states (Table 6), with a global seroprevalence of *T. cruzi* infection of 1.49% [95%CI 1.20-1.85].

**Table 6:** Prevalence of positive children under 18 years by state, 2006-2017 (6 studies).

| State | Sample | Cases | Prevalence (%) | Confidence Interval (95%) |
| --- | --- | --- | --- | --- |
| Estado de México | 356 | 22 | 6.18 | 4.12-9.18 |
| Querétaro | 826 | 11 | 1.33 | 0.74-2.37 |
| Veracruz | 150 | 5 | 3.33 | 1.43-7.56 |
| Yucatán | 700 | 5 | 0.71 | 0.30-1.66 |
| Sub Total | 2,032 | 43 | 2.12 | 1.58-2.84 |
| Without georeference | 3,327 | 37 | 1.11 | 0.81-1.53 |
| Total | 5,359 | 80 | 1.49 | 1.20-1.85 |

\*Cases are reported using 2 or more serological tests to determine the infection.

Several studies also evaluated *T. cruzi* infection in blood donors (9 studies), and positive human cases were detected in every state of the Mexican Republic, except for Baja California Sur, Sinaloa, and Zacatecas (Table 7). The total number of blood donor cases reported was 2,300, corresponding to a national seroprevalence of 0.51% [95%CI 0.49-0.53]. The highest seroprevalence was observed in the states of Quintana Roo, Tabasco, Puebla, Campeche and Nayarit.

**Table 7:** Prevalence of positive serology in blood banks by state, 2006-2017 (9 studies).

| State | Sample | Cases | Prevalence (%) | Confidence Interval (95%) |
| --- | --- | --- | --- | --- |
| Aguascalientes | 7,187 | 1 | 0.01 | 0.00-0.07 |
| Baja California | 6,932 | 16 | 0.23 | 0.14-0.37 |
| Baja California Sur | 375 | 0 | 0 | 0.00-1.01 |
| Campeche | 1,530 | 20 | 1.31 | 0.85-2.01 |
| Chiapas | 3,873 | 12 | 0.31 | 0.18-0.54 |
| Chihuahua | 6,012 | 26 | 0.43 | 0.29-0.63 |
| Coahuila | 4,676 | 12 | 0.26 | 0.15-0.45 |
| Colima | 636 | 5 | 0.79 | 0.34-1.83 |
| Durango | 1,151 | 4 | 0.35 | 0.14-0.89 |
| Estado de México | 93,462 | 209 | 0.22 | 0.19-0.25 |
| Guanajuato | 6,286 | 8 | 0.13 | 0.07-0.25 |
| Guerrero | 4,480 | 20 | 0.45 | 0.29-0.69 |
| Hidalgo | 4,117 | 30 | 0.73 | 0.51-1.04 |
| Jalisco | 12,333 | 55 | 0.45 | 0.35-0.58 |
| México city | 5055 | 15 | 0.30 | 0.18-0.49 |
| Morelos | 3,403 | 26 | 0.76 | 0.52-1.11 |
| Nayarit | 5,307 | 64 | 1.21 | 0.95-1.54 |
| Nuevo León | 37,250 | 100 | 0.27 | 0.22-0.33 |
| Oaxaca | 1,554 | 4 | 0.26 | 0.10-0.66 |
| Puebla | 10,128 | 177 | 1.75 | 1.51-2.02 |
| Querétaro | 3,870 | 22 | 0.59 | 0.38-0.86 |
| Quintana Roo | 2,768 | 55 | 1.99 | 1.53-2.58 |
| San Luis Potosí | 1,532 | 4 | 0.26 | 0.10-0.67 |
| Sinaloa | 1,946 | 0 | 0 | 0.00-0.20 |
| Sonora | 8,947 | 12 | 0.13 | 0.07-0.23 |
| Tabasco | 1,893 | 34 | 1.80 | 1.29-2.50 |
| Tamaulipas | 8,389 | 58 | 0.69 | 0.53-0.89 |
| Tlaxcala | 532 | 3 | 0.56 | 0.19-1.64 |
| Veracruz | 107,251 | 625 | 0.58 | 0.54-0.63 |
| Yucatán | 99,388 | 683 | 0.69 | 0.64-0.74 |
| Zacatecas | 1,190 | 0 | 0 | 0.00-0.32 |
| Total | 453,453 | 2,300 | 0.51 | 0.49-0.53 |

\*Cases are reported using 2 or more serological tests to determine the infection.

Finally, we also examined the type of serological test performed in these studies. The most widely used tests were indirect hemagglutination, followed by Chagatest ELISA from Wiener lab, and immunifluorescence assays (Table 8), which represented 65% of all tests used. Several other commercial ELISA tests were also used (24% of tests), and in-house tests including ELISA, microscopy and western blot represented 5.7% of the tests used.

**Table 8:** Type of diagnostic tests used in epidemiological studies in Mexico

| Type of test | Sample | Cases |
| --- | --- | --- |
| IHA, Indirect hemagglutination | 391,148 | 2,005 |
| Chagatest Elisa (v. 3.0, 4.0 or unspecified), Wiener Lab | 234,282 | 1,234 |
| IFA, Immunofluorescence assays | 103,396 | 962 |
| BioElisa Chagas, Biokit/Werfen | 89,021 | 448 |
| Chagas Recombinant Micro-Elisa Kit, Accutrack | 88,787 | 558 |
| Elisa Kit, Bioschile | 87,342 | 672 |
| In-House Elisa with Antigenic Extract | 34,832 | 521 |
| PCR, Polymerase chain reaction | 25,876 | 75 |
| Microscopy | 24,262 | 19 |
| Chagas STAT-PAK rapid test, Chembio | 18,208 | 293 |
| Abbott Chagas' EIA Kit, Abbott Lab Meridian | 7,296 | 55 |
| Chagas Elisa Kit, Meridian Biosci. | 7,296 | 55 |
| RIPA, Radioimmunoprecipitation | 7,296 | 55 |
| WB, Western blot | 5,379 | 216 |
| Enzyme Chagas Kit, Interbiol | 214 | 8 |
| Chagatek Elisa, Lomos-Biomerieux | 184 | 62 |
| Total | 1,124,819 | 7,238 |

## Discussion

Chagas disease remains one of the most relevant parasitic disease in the Americas, but its epidemiology in Mexico is still poorly understood. Better data are thus urgently needed to help develop appropriate public health programs for disease control and patient care. In this study we analyzed published data on *T. cruzi* seroprevalence of infection in Mexico between 2006 and 2017. A total of 38 studies were identified, covering most of the country with the notable exception of the state of Michoacán. Discrepancies in previous studies have been often attributed to diagnostic methods and uncertainties about the confirmation of cases [57], leading to current recommendations of health agencies requesting a minimum of 2 serological techniques for accurate diagnostic [10]. Based on this criterion, we found a national seroprevalence of *T. cruzi* infection of 2.26%, corresponding to 2.71 million cases in the country. Importantly, few studies were discarded for lack of confirmatory testing, indicating that most of recent studies followed current guidelines for the accurate diagnostic of cases. This seroprevalence level can thus be considered rather conservative, but it is much higher than previous estimates. For example, the Pan American Health Organization (PAHO) estimated that 1,100,000 individuals were infected with *T. cruzi* in Mexico in 2006, and 29,500,000 were at risk of infection [58], and a national prevalence of 0.65% (with 733,333 cases) was established in 2010 by the Mexican Ministry of Health [59]. The most recent estimates from the WHO based on 2010 data reports 876,458 cases (WHO 2015 epidemiological record), corresponding to a national prevalence of 0.78%. Our analysis of data from the last decade thus suggests that the magnitude of *T. cruzi* infection in Mexico may have been underestimated in these previous reports. In addition, recent studies pointing out a low sensitivity of commercial serological tests for *T. cruzi* diagnostic [21,31,60], some of which are well used in Mexico (Table 8) also raise concerns that the seroprevalence of *T. cruzi* infection may even be higher than currently detected. Improvements in serological tests are thus urgently needed for a more reliable disease surveillance [61]. Our analysis nonetheless places Mexico as the country with the largest number of cases of *T. cruzi* infection as previously estimated [10], and highlight the urgency of establishing national priorities for the control of parasite transmission and patient care as well as improved epidemiologic surveillance.

Our results also point out to some regional differences in *T. cruzi* infection seroprevalence among states. The ecology and epidemiology of Chagas disease are the result of many geographical, ecological, biological, and social interactions [62], which may explain some of these differences. High seroprevalence levels have been previously reported for several states including Jalisco, Chiapas, Queretaro, Oaxaca, Veracruz, and Morelos [11], suggesting a well-established endemicity in these states. States with seroprevalence levels higher than previously reported also emerged through our study, in spite of limited sample sizes. These include San Luis Potosi, Estado de Mexico, Hidalgo and Guanajuato.

*T. cruzi* infection is also present at a significant seroprevalence in pregnant women in Mexico. Despite the limited information available for this specific population, we could estimate that there are 22,930 births from *T. cruzi* infected pregnant women per year, corresponding to 1,445 cases of congenitally infected newborns per year in the country. This is very similar to previous estimates [63], suggesting that the epidemiology of congenital Chagas disease has not changed in the past decade. Because infected newborns can be effectively treated, the lack of specific screening programs to identify them is a missed opportunity for the control of the disease. Indeed, a recent health economic study in the US evidenced the large benefits of maternal screening for *T. cruzi* infection, as lifetime societal savings due to screening and treatment was estimated at $634 million saved for every birth year cohort [64].

Very limited information is available on *T. cruzi* infection in children. Nonetheless, we were able to identify a few studies on children up to 18 years, who presented an average prevalence of 1.49%. This may indicate more recent and active transmission compared to data on adult populations, and suggests that the incidence of *T. cruzi* infection has been fairly stable over time. Therefore, effective vector control programs tailored to the extensive diversity of triatomine species present in Mexico [65] are urgently needed to reduce vectorial *T. cruzi* transmission to human populations [66,67].

Blood transfusion has been considered the second most important mode of transmission of Chagas disease in Mexico [68]. In 1998, the screening of almost 65,000 blood donors from 18 government-run transfusion centers showed a 1.5% prevalence of anti-*T. cruzi* antibodies in blood donors [69]. The highest prevalence was detected in the states of Hidalgo, Tlaxcala, Puebla, Chiapas y Yucatan, as expected from previous reports, whereas the northern states of Nuevo Leon and Chihuahua, had the lowest seroprevalence in blood donors. For the period between 1978-2004, Cruz-Reyes *et al.* defined a national prevalence of positive serology in blood banks of 2.03% [11]. In our study, the national prevalence detected in blood donors was lower with 0.51%. These differences can be explained by the increased reliability of serologic screening of blood donors with the passing of legislation making screening mandatory in the year 2000 [13,70]. The highest prevalence of 1.99% is detected in the state of Quintana Roo and the lowest in Baja California Sur, Sinaloa and Zacatecas (with a prevalence of 0%).

## Strengths and limitations

A major strength of our analysis was to consider the reliability of serological testing performed, and to ensure it followed WHO recommendation for confirmation of cases using at least a second test. Hence, our estimates of seroprevalence are robust and conservative. On the other hand, there are some limitations. First, some heterogeneity among study designs and particularly sampling strategies and recruitment of subjects may have generated some bias. Also, publication bias leading to uneven coverage of the different states by research studies may be a confounding factor affecting differences in *T. cruzi* infection prevalence among states, and may also affect the national average, as we did not attempt to weigh studies based on sample sizes and state population. This highlights the need for much improved nationwide disease surveillance to clearly identify geographic heterogeneities in *T. cruzi* transmission and Chagas disease epidemiology. Finally, the small number of studies/sample sizes for some of the subgroup analysis also add uncertainties to our estimates of *T. cruzi* seroprevalence in subpopulations.

## Conclusion

In conclusion, our systematic review shows a national seroprevalence of *T. cruzi* infection of 2.26%, with 2.71 million cases in Mexico, which is higher than previously recognized. It places Mexico as the country with the largest number of cases, highlighting the urgency of establishing Chagas disease control as a key national public health priority, to ensure that it does not remain a major barrier to the economic and social development of Mexico’s most vulnerable populations. The presence of *T. cruzi* infection in specific subpopulations such as pregnant women, children and blood donors also informs on specific risks of infection, and call for the implementation of well-established control interventions [56,67,71]. It also remains essential to strengthen effective nationwide surveillance for Chagas disease, in order to obtain more precise data on its prevalence across the country and its changes over time. Finally, while our estimates are conservative and based on confirmed cases, the lack of sensitivity of current serological tests observed in Mexico suggest that the true magnitude of Chagas disease in the country may still be underestimated, and the development of more reliable diagnostic tests will be key for an effective identification of cases as well as improved patient care [61].

## Acknowledgements

We thank Dr. Pierre Buekens for his helpful comments on a draft version of the manuscript.

## Supporting information

S1: PRISMA checklist

S2: PRISMA flow diagram

